# Distance-based Protein Folding Powered by Deep Learning

**DOI:** 10.1101/465955

**Authors:** Jinbo Xu

## Abstract

Direct coupling analysis (DCA) for protein folding has made very good progress, but it is not effective for proteins that lack many sequence homologs, even coupled with time-consuming folding simulation. We show that we can accurately predict the distance matrix of a protein by deep learning, even for proteins with ∼60 sequence homologs. Using only the geometric constraints given by the resulting distance matrix we may construct 3D models without involving any folding simulation. Our method successfully folded 21 of the 37 CASP12 hard targets with a median family size of 58 effective sequence homologs within 4 hours on a Linux computer of 20 CPUs. In contrast, DCA cannot fold any of these hard targets in the absence of folding simulation, and the best CASP12 group folded only 11 of them by integrating DCA-predicted contacts into complex, fragment-based folding simulation. Rigorous experimental validation in CASP13 shows that our distance-based folding server successfully folded 17 of 32 hard targets (with a median family size of 36 sequence homologs) and obtained 70% precision on top L/5 long-range predicted contacts. Latest experimental validation in CAMEO shows that our server predicted correct fold for two membrane proteins of new fold while all the other servers failed. These results imply that it is now feasible to predict correct fold for proteins lack of similar structures in PDB on a personal computer without folding simulation.

**Significance:** Accurate description of protein structure and function is a fundamental step towards understanding biological life and highly relevant in the development of therapeutics. Although greatly improved, experimental protein structure determination is still low-throughput and costly, especially for membrane proteins. As such, computational structure prediction is often resorted. Predicting the structure of a protein with a new fold (i.e., without similar structures in PDB) is very challenging and usually needs a large amount of computing power. This paper shows that by using a powerful deep learning technique, even with only a personal computer we can predict new folds much more accurately than ever before. This method also works well on membrane protein folding.

## Introduction

De novo protein structure prediction from sequence alone is one of the most challenging problems in computational biology. Even after decades of research, progress on this problem was slow, and many methods require considerable computational resources, even for relatively small proteins. Nevertheless, in recent years good progress has been achieved thanks to accurate contact prediction enabled by direct coupling analysis (DCA)^1-9^ and deep convolutional neural networks (DCNN)^10-16^. As such, contact-assisted protein folding has gained a lot of attention and contact prediction has garnered considerable research effort.

We have developed the CASP12-and CASP13-winning method RaptorX-Contact^10^ that uses deep and global convolutional residual neural network (ResNet) to predict contacts. ResNet is one type of DCNN^17^, but much more powerful than traditional or plain DCNN. RaptorX-Contact has good accuracy even for some proteins with only dozens of effective sequence homologs. The accuracy of RaptorX-Contact decreases much more slowly than DCA when more predicted contacts are evaluated even when the protein under study has thousands of sequence homologs (see Table 1 in the RaptorX-Contact paper^10^). As reported in^10, 12^, without folding simulation, the 3D models constructed from contacts predicted by RaptorX-Contact have much better accuracy than those built from contacts predicted by DCA methods such as CCMpred^6^ and the CASP11 winner MetaPSICOV^18^ (equivalent to a feed-forward neural network of 3 hidden layers). RaptorX-Contact also works well for membrane proteins even trained by soluble proteins^12^ and for complex contact prediction even trained by single-chain proteins^19^.

**Table 1.** The average quality (TMscore) of the 3D models predicted by various methods for the 37 CASP12 hard targets. Top k (k=1, 5) means that for each target the best of top k 3D models is considered. Columns “Top 1” and “Top 5” list the average TMscore of the top 1 and the best of top 5 models for all the targets, respectively. Column “#correct” lists the number of targets with correctly predicted folds (TMscore>0.5) when all 5 models are considered. The results in rows 3-9 are copied from our previous paper^11^.

| Group | Top 1 | Top 5 | #correct (TM>0.5) |
| --- | --- | --- | --- |
| **This work** | 0.466 | 0.476 | 21 |
| Our contact | 0.354 | 0.397 | 10 |
| CCMpred | 0.216 | 0.235 | 0 |
| MetaPSICOV | 0.262 | 0.289 | 2 |
| Baker-server | 0.326 | 0.370 | 9 |
| Zhang-server | 0.347 | 0.404 | 10 |
| Baker-human | 0.392 | 0.422 | 11 |
| Zhang-human | 0.375 | 0.420 | 11 |

Both (global) ResNet and DCA are global prediction methods because they predict whether one pair of residues form a contact by considering the status of the other residue pairs, which is the key to the significant improvement in contact prediction. However, ResNet is better than DCA in that 1) ResNet can capture higher-order residue correlation (i.e., structure motifs) while DCA mainly focuses on pairwise relationship; 2) ResNet tries to learn the global context of a contact matrix and uses this information to predict the status of one residue pair; and 3) existing DCA methods are roughly linear models with tens of millions of parameters to be estimated from a single protein family, while ResNet is a nonlinear model with fewer parameters to be estimated from thousands of protein families. Before this (global) ResNet method was proposed for protein folding, deep learning (DL) models such as CMAPpro^20^ and Deep Belief Networks (DBN)^21^ were used for contact prediction, but ResNet is the first DL method that greatly outperforms DCA and shallow methods such as MetaPSICOV^18^. Different from (global) ResNet and DCA, DBN and MetaPSICOV are local prediction methods, as these methods predict the status of one residue pair without considering the status of other residue pairs. This is one of the major reasons why RaptorX-Contact greatly outperformed DBN and MetaPSICOV. Inspired by the success of RaptorX-Contact, most CASP13 contact predictors have employed global ResNet or DCNN^13, 15, 22^, as shown in <u>the CASP13 abstract book</u>. Notably, Cheng group, who developed DBN, has switched to DCNN for contact prediction^13^. Peng group, who employed traditional DCNN in CASP12^16^, has switched to ResNet in CASP13.

Though contact prediction is drawing considerable attention, in this paper we study distance prediction and treat contact prediction as a by-product. The rationale for this decision is two-fold. The distance matrix contains finer-grained information than contact matrix and provides more physical constraints of a protein structure, e.g., distance is metric while contact is not. Because of this, a distance matrix can determine a protein structure (except mirror image) much more accurately than a contact matrix. Trained by distance instead of contact matrices, ResNet may automatically learn more about the intrinsic properties of a protein structure and thus, greatly reduce the conformation space and improve folding accuracy. Further, different from DCA that aims to predict only a small number of contacts and then use them to assist folding simulation, we would like to predict the whole distance matrix and then directly construct protein 3D models without invoking any folding simulation at all. By doing this, we significantly reduce running time needed for protein folding, especially for a large protein. As we use many more distance restraints to construct 3D models, the impact of individual distance prediction error may be reduced (by the law of large numbers). In contrast, contact-assisted folding simulation may be misguided by several wrongly-predicted contacts and needs a long time to generate a good conformation for a large protein.

Although there are few studies on distance prediction, it is not totally new. For example, Aszodi et al^23^ predicted inter-residue distance from pairwise conserved hydrophobicity score and used it to reconstruct 3D models of a protein. Kloczkowski et al^24^ described a spectral decomposition method for inter-residue distance prediction. Pietal et al.^25^ predicted the inter-residue distance matrix from a contact matrix and then built protein 3D models from predicted distance. Kukic et al.^26^ proposed a recursive neural network method for inter-residue distance prediction and studied distance-based protein folding, but the predicted distance has a large error and the resultant 3D models have poor quality. In 2012, we employed a probabilistic neural network to predict inter-residue distance and then derived protein-and position-specific statistical potential from predicted distance distribution^27^. We have also studied folding simulation using this distance-based statistical potential^28^. Recently, we showed that protein-specific distance potential derived from deep ResNet may improve by a large margin^29^ protein threading with weakly similar templates.

In addition to inter-atom distance prediction, we also employ deep ResNet to predict secondary structure and backbone torsion angles, although these features are much less important than distance for folding. By feeding these three types of predicted restraints to CNS^30^, a computer program for experimental protein structure determination that does not use any folding simulation, we are able to construct more accurate 3D models than what can be achieved from predicted contacts, as evidenced by our experiments on a set of 37 CASP12 hard targets^14^ and a set of 41 CAMEO hard targets^10^. Our distance-based ab initio folding method predicts many more correct folds for the CASP12 targets than the best CASP12 group that integrated contacts, fragments, server predictions and conformation sampling into a complex folding simulation protocol. Rigorous experimental validation in CASP13 shows that our distance-based folding server achieved the best contact prediction accuracy among 46 groups and successfully folded 17 of 32 hard targets. Latest rigorous experimental validation in CAMEO shows that our ab initio folding server predicted correct fold for two membrane proteins while all the other CAMEO-participating servers failed. Finally, our distance-based folding algorithm runs very fast, folding the 41 CAMEO targets within 13 hours and the 37 CASP12 targets within 4 hours on a single Linux computer of 20 CPUs, excluding time for feature generation.

## Results

### Results on the 37 CASP12 hard targets

Table 1 summarizes folding accuracy of various methods on the 37 CASP12 hard targets, including our distance-based ab initio folding method, three contact-based ab initio folding methods (i.e., our own method, CCMpred and MetaPSICOV) and 4 top CASP12 groups (Baker-server^39^, Baker-human^40^, Zhang-server^41^ and Zhang-human^42^). As shown in Table 1, the 3D models predicted by our distance-based ab initio folding method have average TMscore 0.466 and 0.476, respectively, when the top 1 and the best of top 5 models are evaluated. These scores are much better than the models built from contacts predicted by ourselves, CCMpred and MetaPSICOV. Our distance-based 3D models also have much higher quality than the 4 top CASP12 groups, which folded proteins by combining fragment assembly, predicted contacts, conformation sampling and Monte Carlo search. Zhang-human also extracted information from the CASP12 server predictions. By contrast, our folding algorithm does not use any template fragments or any CASP12 server predictions.

When all 5 models are considered, our distance-based folding can predict correct folds (TMscore>0.5) for 21 of the 37 targets while our contact-based folding can do so for only 9 of them and the best CASP12 human group can do so for only 11 of them. See Figure 1(A) for the detailed comparison between our distance-based and contact-based models. With only predicted contacts and predicted secondary structure, MetaPSICOV generates correct fold for only 2 targets and CCMpred cannot generate any correct folds. That is, the contacts predicted by CCMpred and MetaPSICOV alone are not good enough for 3D modeling.

**Figure 1.**
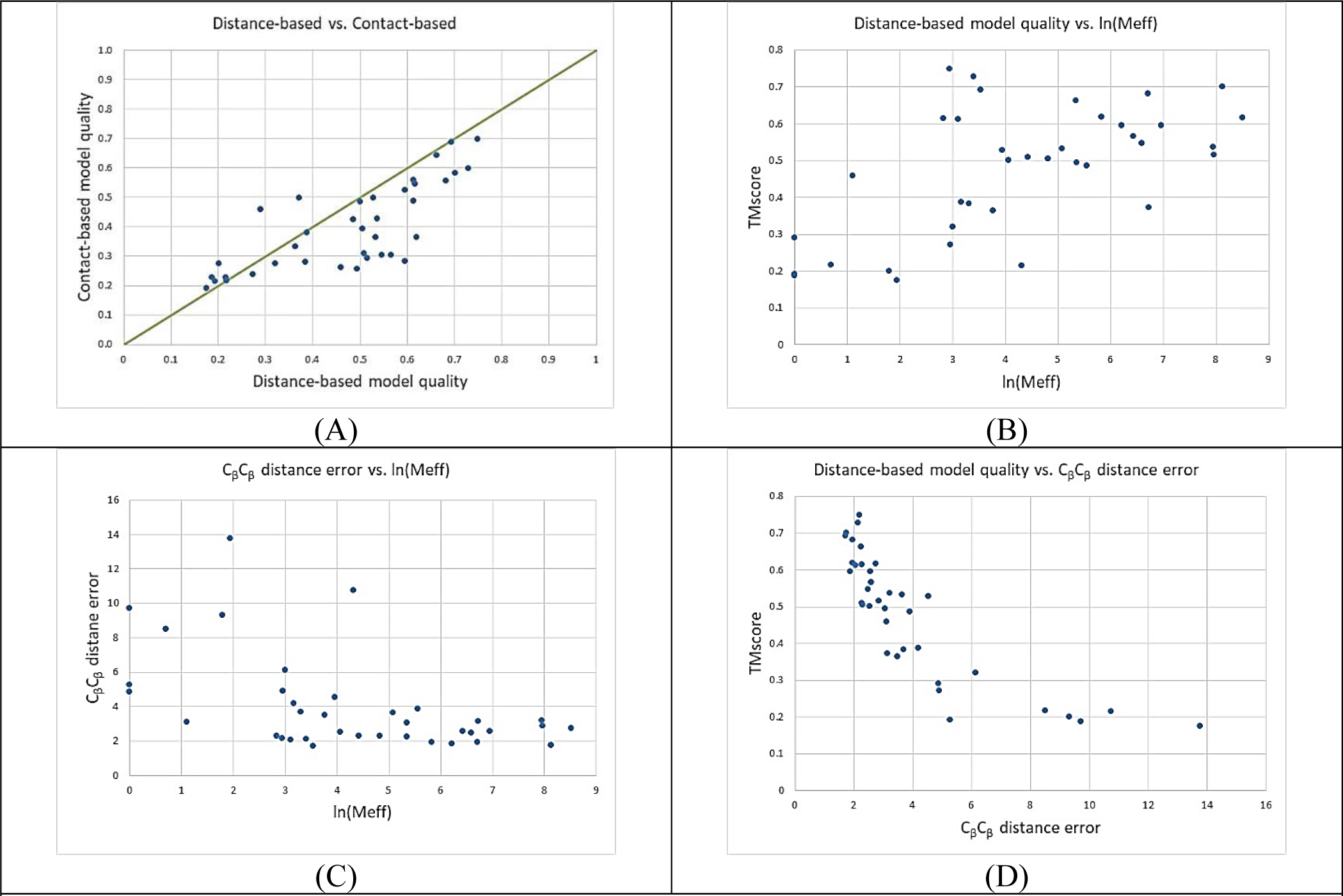
Results on the 37 CASP12 hard targets. (A) distance-based 3D model quality vs. contact-based 3D model quality; (B) distance-based 3D model quality vs. Meff; (C) C_β_-C_β_ distance prediction error vs. Meff. (D) distance-based 3D model quality vs. C_β_-C_β_ distance prediction error.

Figure 1(B) shows that the quality of our distance-based 3D models is correlated to Meff, which measures the number of sequence homologs of a target. When Meff>55 or ln(Meff)>4, there is a good chance that our predicted 3D models have a correct fold. Our distance-based ab initio folding can fold 8 out of 21 targets with Meff≤100: T0862-D1, T0863-D1, T0869-D1, T0870-D1, T0894-D1, T0898-D1, T0904-D1 and T0915-D1. Meanwhile, 5 of them have 3D models with TMscore>0.6. In contrast, Zhang-Server, Zhang-Human, Baker-Server and Baker-Human predicted models with TMscore>0.6 for only 2, 1, 0 and 0 targets with Meff≤100, respectively.

#### Evaluation of distance prediction

Here we only consider the pairs of atoms with sequence separation at least 12 residues and predicted distance≤15Å. Table 2 summarizes the quality of predicted distance on the 37 hard CASP12 targets. The quality is first calculated on each target and then averaged across all the test targets. Figure 1(C) shows that there is certain correlation between distance prediction error and Meff. Figure 1(D) shows a strong correlation between 3D modeling quality and distance prediction error, which means that as long as distance prediction is accurate, CNS is able to build good 3D models. When the distance error is 8Å, the predicted 3D models are equivalent to a random model in terms of quality. By the way, the top L, L/2, L/5 and L/10 long-range contact accuracy is 43.1%, 56.9%, 66.8% and 73.7%, respectively, on average ∼6% better than what we have reported before^11^. Note that here we still used the multiple sequence alignments (MSA) generated in 2016 for the 37 targets.

**Table 2.** Quality of predicted distance on the 37 hard CASP12 targets.

| Atom pair | Absolute error | Relative error | Precision | Recall | F1 |
| --- | --- | --- | --- | --- | --- |
| C<sub>β</sub>C<sub>β</sub> | 4.045Å | 0.269 | 0.6541 | 0.5106 | 0.5677 |
| C<sub>g</sub>C<sub>g</sub> | 4.498Å | 0.296 | 0.6123 | 0.5153 | 0.5529 |
| C<sub>α</sub>C<sub>g</sub> | 4.421Å | 0.285 | 0.6323 | 0.4955 | 0.5485 |
| C<sub>α</sub>C<sub>α</sub> | 3.938Å | 0.257 | 0.6499 | 0.5229 | 0.5733 |
| NO | 4.035Å | 0.262 | 0.6412 | 0.4996 | 0.5563 |

#### Importance of Direct Information (DI)

DI produced by DCA is an important input feature for RaptorX-Contact. To evaluate its importance for contact prediction and folding, we trained one ResNet model without using DI, but keeping other information including mutual information (MI). This ResNet model has top L, L/2, L/5 and L/10 long-range contact prediction accuracy 36.3%, 48.4%, 61.9% and 66.7%, respectively. The average TMscores of the top 1 and best of top 5 3D models for the 37 targets are 0.400 and 0.411, respectively. That is, the long-range contact prediction accuracy drops by ∼7% and the 3D model quality drops by 0.066 TMscore when DI is not used. The performance decrease is not very large because many of the 37 targets have a small number of sequence homologs and thus, DI is not much better than MI.

### Results on the 41 CAMEO hard targets

On average, this dataset is easier than the CASP12 set. Again, we build 3D models using CNS from the predicted distance and angle restraints. Our folding algorithm does not use any template fragments or predictions produced by any other servers. In summary, the results on this dataset are consistent with that on the CASP12 test set. When the first models and the best of top 5 models are evaluated, our distance-based ab initio folding algorithm has average TMscore 0.551 and 0.577, respectively, about 10% better than our contact-based ab initio folding algorithm, which has average TMscore 0.504 and 0.524, respectively. Figure 2(A) shows that for most targets, our distance-based models have better quality than our contact-based models. In total, our distance- and contact-based methods predicted correct folds for 30 and 23 targets, respectively. Figure 2(B) shows that our distance-based model quality is correlated to Meff (i.e., the number of sequence homologs available for a target), but our method predicted a correct fold for 1 target with Meff=1. Similar to the observation on the CASP12 dataset, when Meff>55 or ln(Meff)>4, there is a good chance that our predicted 3D models have a correct fold. Figure 2(D) shows that there is strong correlation between 3D modeling accuracy and distance prediction error. Finally, when all top 5 models are considered, the 3D models generated by CCMpred contacts and MetaPSICOV contacts have average TMscore 0.316 and 0.392, respectively, much worse than our distance-based method.

**Figure 2.**
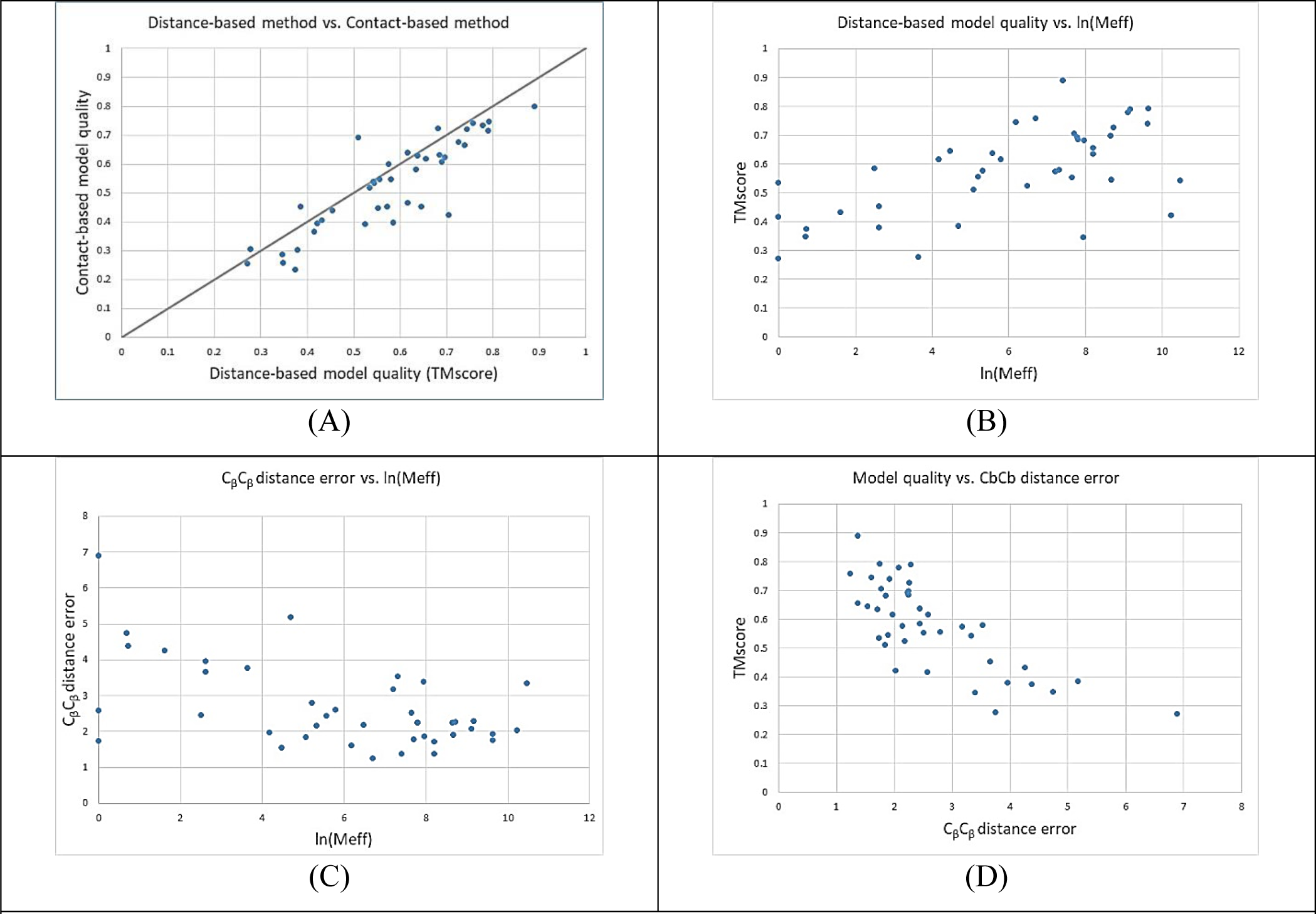
Results on the 41 CAMEO hard targets. (A) distance-based model quality vs. contact-based model quality; (B) distance-based 3D model quality vs. Meff; (C) C_β_-C_β_ distance prediction error vs. Meff; (D) distance-based 3D model quality vs. C_β_-C_β_ distance prediction error.

#### Evaluation of distance prediction

Here we only consider the pairs of atoms with sequence separation at least 12 residues and predicted distance≤15Å. Table 3 summarizes the average quality of predicted distance on the 41 CAMEO hard targets. Figure 2(C) shows that the distance prediction error is inversely proportional to the logarithm of Meff. When Meff>55 or ln(Meff)>4, there is a very good chance that the average C_β_-C_β_ distance prediction error is less than 4Å. This is consistent with what is shown in Figure 1(C).

**Table 3.** Quality of predicted distance on the 41 CAMEO hard targets.

| Atom pair | Absolute error | Relative error | Precision | Recall | F1 |
| --- | --- | --- | --- | --- | --- |
| C<sub>β</sub>C<sub>β</sub> | 2.725Å | 0.207 | 0.6941 | 0.6133 | 0.6158 |
| C<sub>g</sub>C<sub>g</sub> | 3.212Å | 0.237 | 0.6580 | 0.6074 | 0.5921 |
| C<sub>α</sub>C<sub>g</sub> | 3.058Å | 0.223 | 0.6687 | 0.5653 | 0.5914 |
| C<sub>α</sub>C<sub>α</sub> | 2.726Å | 0.197 | 0.7032 | 0.5943 | 0.6230 |
| NO | 2.814Å | 0.196 | 0.6987 | 0.5770 | 0.6139 |

### Rigorous experimental validation in CASP13

We have tested our distance-based folding algorithm in the rigorous blind test CASP13 in 2018, which in total has 32 hard targets for servers (and 31 for human groups). We registered RaptorX-Contact for contact prediction and distance-based ab initio folding, and RaptorX-DeepModeller for distance-based folding of a target with only weakly similar templates. RaptorX-DeepModeller differs from RaptorX-Contact in that the ResNet model used for RaptorX-DeepModeller has one additional input feature, namely, an initial distance matrix extracted from the weakly similar template according to the alignment. Supposing two target residues i and j are aligned to two template residues k and l, we assign the distance between k and l as the initial distance of i and j. When one target residue is not aligned, the corresponding row and column in the initial distance matrix is empty.

RaptorX-Contact was officially ranked first among 46 (human and server) contact prediction groups. The CASP13 assessor evaluated the contact predictors by many different metrics. Here we only report precision and F1 value calculated on top L/5 predicted long-range contacts where L is the sequence length. In terms of precision, the top 5 servers are: RaptorX-Contact (70.054%), TripletRes (65.678%), ResTriplet (64.031%), DMP (60.798%) and TripletRes_AT (60.595%). In terms of F1 value, the top 5 servers are: RaptorX-Contact (0.23276), TripletRes (0.21258), ResTriplet (0.20826), RRMD (0.19176) and TripletRes_AT (0.19126). To the best of our knowledge, all the top contact prediction groups used deep ResNet. As a control, MetaPSICOV ran by the CASP13 organizers has precision and F1-value 25.16% and 0.0784, respectively and a DCA method GaussDCA has precision and F1-value 21.757% and 0.06732, respectively. Table 4 shows the average quality of our predicted distance on the CASP13 hard targets, which is slightly better than that of the CASP12 hard targets.

**Table 4.** Quality of predicted distance on the 32 CASP13 hard targets.

| Atom Pair | Absolute error | Relative error | Precision | Recall | F1 |
| --- | --- | --- | --- | --- | --- |
| CbCb | 3.762Å | 0.2585 | 0.6779 | 0.5397 | 0.5877 |
| CgCg | 4.016Å | 0.2776 | 0.6562 | 0.5315 | 0.5725 |
| CaCg | 3.838Å | 0.2616 | 0.6707 | 0.5117 | 0.5668 |
| CaCa | 3.836Å | 0.2530 | 0.6661 | 0.5319 | 0.5766 |
| NO | 3.750Å | 0.2525 | 0.6739 | 0.5050 | 0.5658 |

Our distance-based folding method did well in predicting 3D models for 32 CASP13 hard targets. Table 5 shows the performance of top 5 CASP13 servers when the best models are evaluated. Our two distance-based folding servers are only slightly worse than Zhang’s two servers, which used ResNet-predicted contacts to guide time-consuming folding simulation. If RaptorX-DeepModeller and RaptorX-Contact are merged into a single group, then the best models produced by us have an average TMscore of 0.5264. Similarly, if Zhang-server and QUARK are merged into a single group, the best models produced by Zhang group have an average TMscore of 0.5348. Robetta underperformed the top 4 servers by a large margin since it used the inaccurate DCA-predicted contacts to guide folding simulation. The TMscore and GDT in Table 5 are calculated by ourselves, so they may be slightly different than what are shown in the CASP13 web site.

**Table 5.**
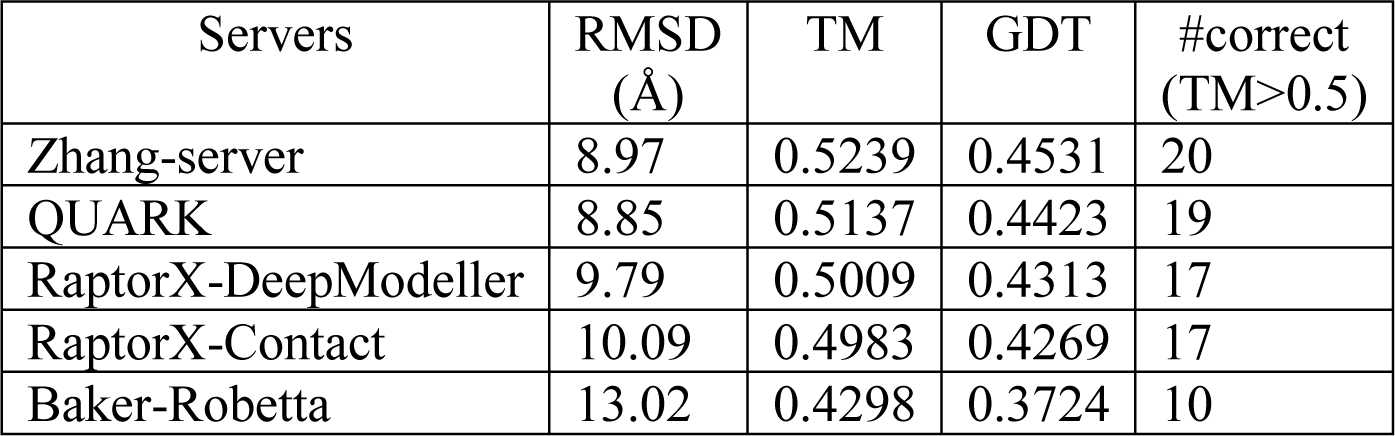
Folding results on 32 CASP13 hard targets when the best of top 5 models is evaluated.

As shown in Fig. 3, there is a good correlation (0.68) between the 3D model quality (TMscore) and the top L/2 long-and medium-range precision. Similar to what we have observed on the CASP12 data, there is also a good correlation between 3D model quality and distance prediction error and between 3D model quality and the logarithm of Meff (i.e., MSA depth). Here we use MSA generated by HHblits and reported in the CASP13 contact assessment page (http://predictioncenter.org/casp13/rrc_results.cgi) so that all the groups may use the same set of Meff values for comparison. Similar to the results on the CASP12 data, when Meff>55 or ln(Meff)>4, there is a good chance that RaptorX-Contact can predict a correct fold (TMscore>0.5). When ln(Meff) is between 3 and 4, RaptorX-Contact may predict correct folds for half of the targets.

**Figure 3.**
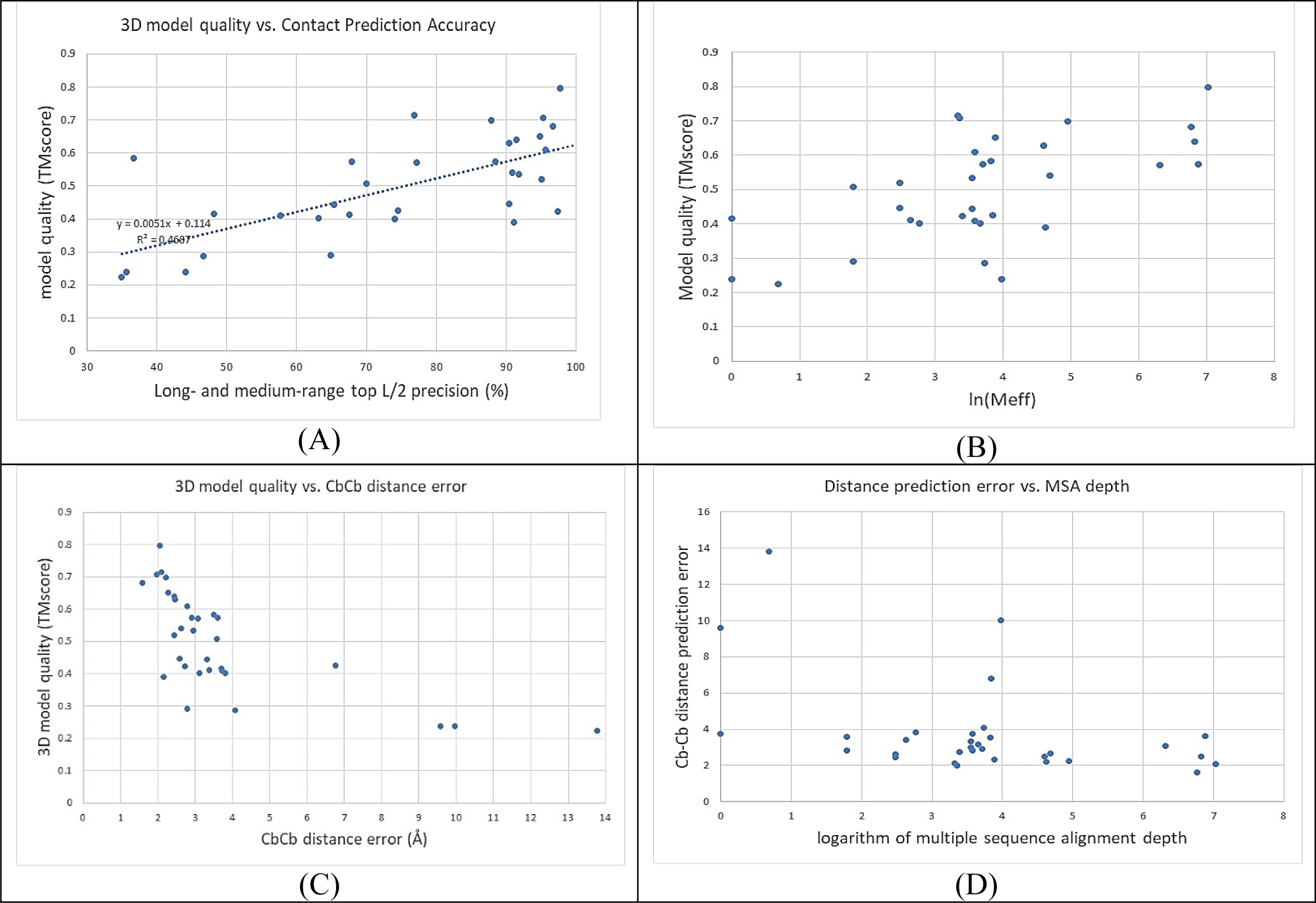
Results of RaptorX-Contact on the 32 CASP13 hard targets. (A) 3D model quality vs. contact prediction accuracy; (B) 3D model quality vs. the logarithm of Meff; (C) 3D model quality vs. C_β_-C_β_ distance prediction error; (D) C_β_-C_β_ distance prediction error vs. the logarithm of Meff.

Among the 32 hard targets, RaptorX-Contact predicted very good 3D models for T0953s2-D2 and T0957s2-D1, both of which have ∼30 sequences in their MSAs. Meanwhile, T0953s2-D2 has 111 residues with valid 3D coordinates and T0957s2-D1 has 155 residues with valid 3D coordinates. Fig. 4 shows the superimposition between their predicted 3D models (blue) and their native structures (red). For both targets, our predicted 3D models have TMscore>0.7 and RMSD 3-4Å. RaptorX-Contact also predicted a very good 3D model (TMscore=0.8, RMSD=5.1Å, picture not shown to save space) for T0969-D1, a α+β protein domain of 354 residues.

**Figure 4.**
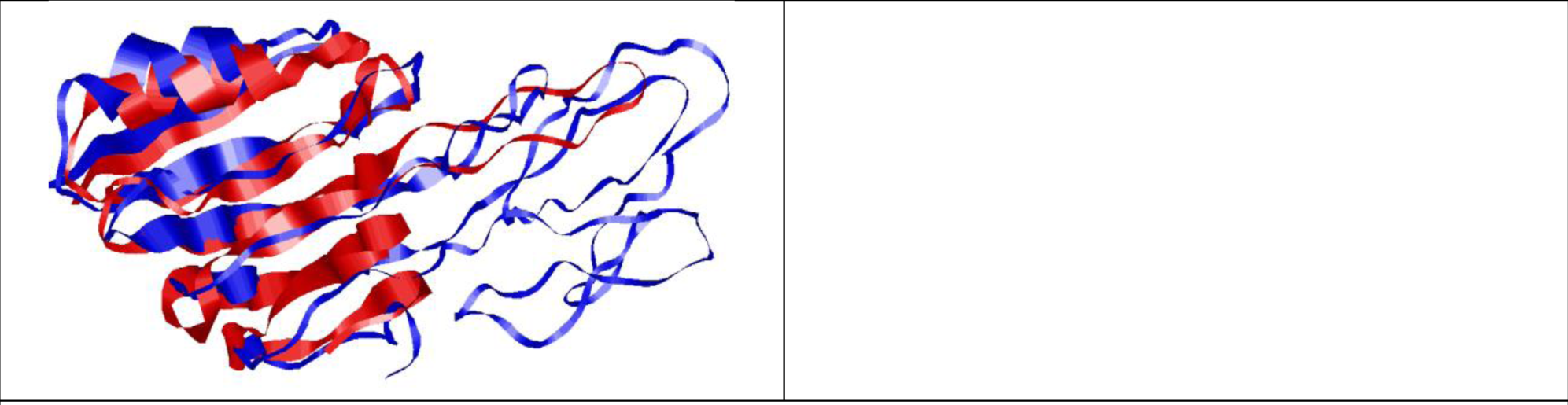
Superimposition between 3D models (blue) predicted by RaptorX-Contact and native structure (red). Left: T0953s2-D2, RMSD=3.33Å, TMscore=0.73; Right: T0957s2-D1, RMSD=4.23Å, TMscore=0.713. The TMscore and RMSD are taken from the CASP13 web site

### Rigorous experimental validation in CAMEO

In mid-September 2018, we started to test our distance-based folding server in the online benchmark CAMEO (http://www.cameo3d.org/) operated by Schwede group. CAMEO is a blind test and the native structure of a test protein is not available until one week after the test. Currently, CAMEO is testing ∼40 web servers including some popular ones such as Robetta, Phyre, RaptorX, Swiss-Model, IntFold and HHpred. Here we only report the result of two recent targets (CAMEO-3D IDs: 2018-11-03_00000053_1 and 2018-11-17_00000062_1) since they are membrane proteins of new fold and CASP13 does not seem to have this kind of targets. The first target has a PDB ID 6bhp, chain name C, 200 residues and 229 effective sequence homologs (based upon the MSA generated by HHblits). As shown in Fig. 5, our ab initio folding server RaptorX-Contact (ID: server 60) predicted 3D models with TMscore=0.68 and RMSD=5.65Å while the other servers failed to predict a correct fold. Further, our response time is much shorter than Baker’s Robetta. Meanwhile, this job was queued in our server for about 4.5 hours and the actual time spent in folding this target is about 1.5 hours. The second target has a PDB ID 6a2j, chain name A, 309 residues and a few thousand sequence homologs. RaptorX-Contact predicted 3D models (picture not shown to save space) with TMscore=0.73 and RMSD=5.51Å, while the other servers predicted 3D models with TMscore≤0.4.

**Figure 5.**
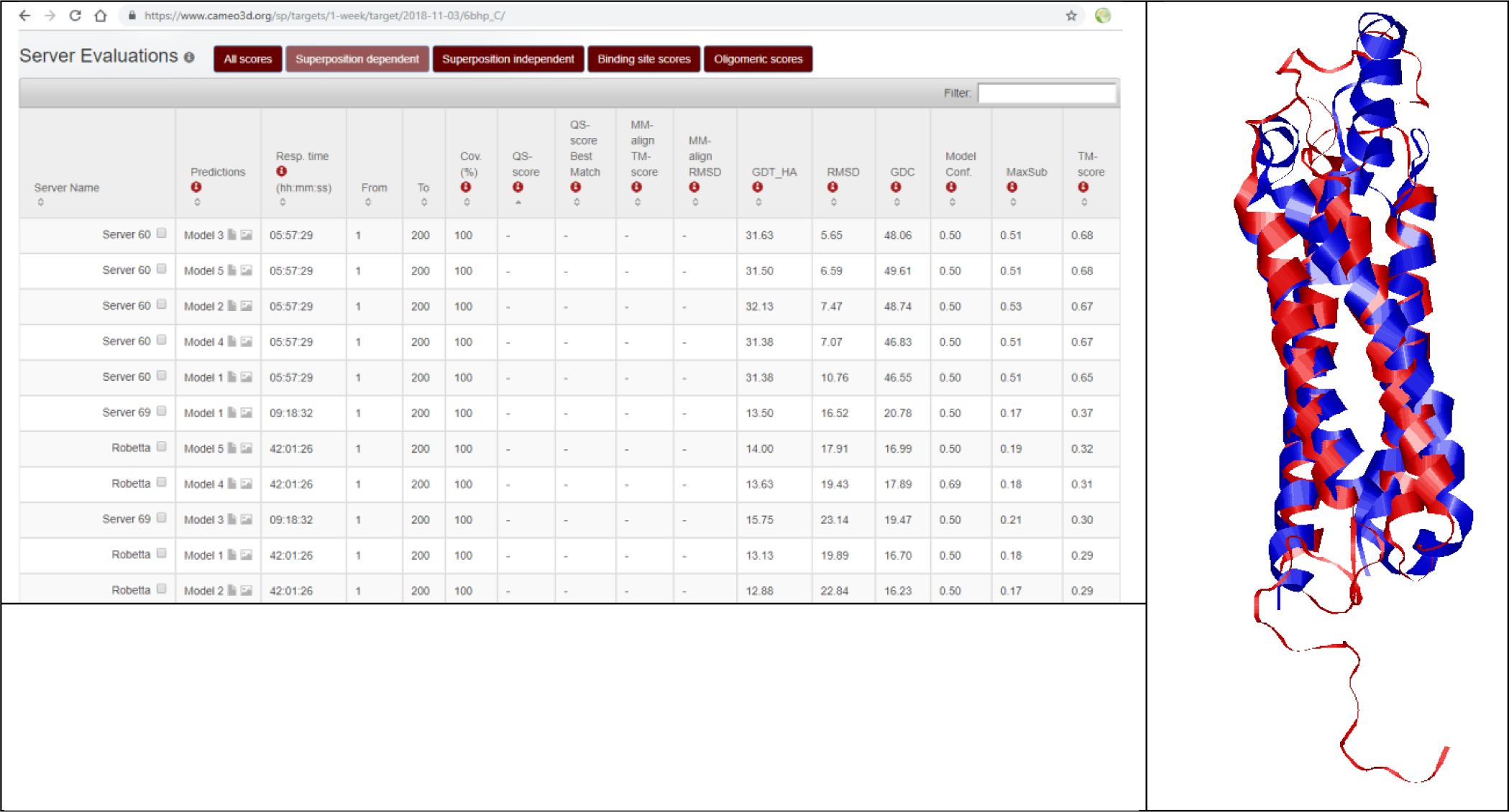
(Top) A screenshot showing the quality of the top 3D models submitted to CAMEO for a target (ID: 2018-11-03_00000053_1) ranked in decreasing order of TMscore. (Right) Superimposition between a 3D model (red) predicted by RaptorX-Contact and native.

### Running time

Our distance-based ab initio folding algorithm runs very fast since it does not involve folding simulation. The whole folding pipeline consists of three main steps: 1) generating multiple sequence alignments and input features; 2) predicting angles and distance; and 3) folding by feeding distance and angle restraints to CNS. It takes minutes to finish the first step for most targets, seconds to finish the second step on a GPU card, and 10 minutes to a few hours to finish the third step on a Linux computer of 20 CPUs. By running the third step in parallel on a Linux computer of 20 CPUs, it took in total ∼13 hours to fold the 41 CAMEO targets and ∼4 hours to fold the 37 CASP12 targets.

### Conclusion and Discussion

We have shown that we can predict an inter-atom distance matrix very well and that even without folding simulation, predicted distance can be used to fold many more proteins than ever before. Although our algorithm does not use fragment assembly, complex energy functions or time-consuming folding simulation, it can predict correct folds for many more hard targets than the best CASP12 groups. Rigorous experimental validation in CASP13 and CAMEO confirmed that our distance-based folding algorithm works well for hard targets. Since it does not need folding simulation at all, our algorithm runs very fast, taking from 10 minutes to a few hours to generate 200 decoys on a Linux computer of 20 CPUs. That is, it is now feasible to do ab initio folding on a personal computer equipped with a GPU card.

We do not give a detailed evaluation of the accuracy of our 1D deep residual neural network models for secondary structure and torsion angle prediction because 1) although the methods are different, the prediction accuracy is similar to what we have reported before^43, 44^ and 2) secondary structure and torsion angles are much less important than distance for protein folding. Without using predicted torsion angles, on average, the 3D model quality decreases by ∼0.008 in terms of TMscore. Nevertheless, the predicted torsion angles may help reduce the number of mirror images generated by CNS.

In this study we only reported the folding results when all the 5 types of atom pairs are used. In fact, using only C_β_-C_β_, our distance-based ab initio folding algorithm can generate slightly worse 3D models than using all 5 types of atom pairs because their distance is highly correlated. Among the 5 types of atom pairs, C_β_-C_β_ is the most useful when only the Cα conformation is evaluated. Nevertheless, using all 5 types of atom pairs can help reduce noise a little bit and may improve side chain packing.

We predicted inter-residue distance by discretizing it into 25 bins. We have also experimented with discretizing the distance into 12 bins (i.e., bin width=1Å) and 52 bins (i.e., bin width=0.25Å). On average, there is no big difference between using 25 bins and 52 bins, both of which are better than using 12 bins. Instead of using a discrete representation of distance, we may predict a real-valued distance matrix by assuming that distance has a log-normal distribution^45^ and then revise our deep learning model to predict the mean and variance of the distribution. CNS can easily take the predicted mean and variance as distance restraints to build 3D models. Currently we use only distance≤15Å for 3D model construction. In the future, we will study whether 3D modeling accuracy can be further improved by using the whole real-valued distance matrix, especially for the determination of domain orientation of a multi-domain protein.

Recently an end-to-end deep learning method was proposed by Dr. AlQuraishi that can directly predict 3D coordinates of a protein structure from its sequence profile^46^. The idea is unique and attractive, but a few rigorous test results are needed to show that this idea is indeed effective on hard targets with very few sequence homologs. In CASP13, a human group AlphaFold developed by DeepMind did much better than us. AlphaFold also employed deep ResNet to predict inter-residue distance distribution (which is similar to our work) and then converted predicted distance probability into statistical potential for energy minimization. AlphaFold did better than us mainly because that AlphaFold used Rosetta to do energy minimization and refinement while we did not do so at all. The energy function minimized by AlphaFold is composed of its own distance-based statistical potential and Rosetta’s physics-based energy terms. We are not able to compare us with AlphaFold in terms of contact and distance prediction accuracy since the DeepMind team did not report them. We have sent an email to the team to request this kind of information, but so far have not received any reply. Nevertheless, the results achieved by AlphaFold further confirm that deep ResNet, which was first developed by us for contact/distance prediction, is a powerful method for protein folding and that accurate distance prediction enables us to fold proteins without time-consuming conformation sampling.

## Methods

### Datasets and Methods to Compare

We self-test our distance-based folding methods on 2 datasets: the set of 37 CASP12 hard targets^10, 11^ and the set of 41 CAMEO hard targets^10^. Both datasets have been used to test our contact prediction algorithm before. CASP is a prestigious blind test of protein structure prediction organized by a human committee and CAMEO is an online blind test of structure prediction run by Schwede group^31^. The CASP12 hard targets were released in Summer 2016 and the CAMEO hard targets were collected in September and October 2016. Please see our papers^10, 11^ for their detailed description. To warrant a fair comparison, we did not regenerate the features for the test proteins. Instead we used the same input features for them as described in^10, 11^. We trained and validated our deep learning models using in total ∼10,000 non-redundant proteins deposited to PDB before May 2016 when CASP12 started. This guarantees that the test targets have no similar proteins in the training set. Meanwhile, 600 proteins were randomly selected to form a validation set and the remaining ∼9400 to form a training set. We used 25% sequence identity as the cutoff to determine if two proteins are redundant or not.

For the CASP12 dataset, we compared our method with CCMpred, MetaPSICOV and 4 top CASP12 groups. For the CAMEO dataset, we compared our method with CCMpred and MetaPSICOV. CCMpred is one of the best DCA (direct coupling analysis) methods, as reported in^10, 12^. MetaPSICOV is the best method for contact prediction in CASP11. In addition, we blindly tested our algorithm as server groups in CASP13 in Summer 2018, which in total has 32 free-modeling (i.e., hard) targets. CASP has two types of participants: server groups and human groups. Server groups have only 3 days for a target while human groups have 3 weeks. Human groups can also make use of all server predictions, so we mainly compare our servers with the other CASP13-participating servers.

### MSA (multiple sequence alignment) and protein feature generation

For the CASP12 and CAMEO datasets, to ensure fair comparison with the results in^10, 11^ and the CASP12 groups, we used the same MSAs and protein features as described in^10, 11^ for the test proteins and the same MSAs and protein features as described in^10^ for the training proteins. That is, for each test protein we generated four MSAs by running HHblits with 3 iterations and E-value set to 0.001 and 1, respectively, to search the uniprot20 library released in November 2015 and February 2016, respectively. Since the sequence databases were created before CASP12 started in May 2016, the comparison with the CASP12 groups is fair. For CASP13 targets, we generated their MSAs (and other sequence features) using the UniClust30 library^32^ created in October 2017 and the UniRef sequence database^33^ created early in 2018. From each individual MSA, we derived both sequential and pairwise features. Sequential features include sequence profile and secondary structure as well as solvent accessibility predicted by RaptorX-Property^34^. Pairwise features include mutual information (MI), pairwise contact potential and direct information (DI) generated by CCMpred^6^. In summary, one test protein has 4 sets of input features and accordingly 4 predicted distance matrices, which are then averaged to obtain the final prediction.

### Predict inter-atom distance by deep residual networks (ResNet)

We use a very similar deep learning (DL) method as described in^10^ to predict the Euclidean distance distribution of two atoms (of different residues) in a protein to be folded. The DL model in^10^ is designed for contact prediction (i.e., binary classification) and consists of one 1D deep ResNet, one 2D deep ResNet and one logistic regression (see Fig. S1 in Appendix for a picture). The 1D ResNet is used to capture sequential context of one residue (or sequence motifs) while the 2D network is used to capture pairwise context of a residue pair (or structure motifs). For distance prediction, we replace the traditional convolutional operation in 2D ResNet by a dilated convolutional operation^35^, which is slightly better since it needs fewer model parameters to have the same receptive field.

We discretize inter-atom distance into 25 bins: <4Å, 4-4.5Å, 4.5-5Å, 5-5.5Å, …, 15-15.5Å, 15.5-16Å, and >16Å. That is, we use 25 labels for distance prediction, as opposed to 2 labels for contact prediction. The DL model for distance prediction is trained using the same procedure as described in^10^. By summing up the predicted probability values of the first 9 distance labels (corresponding to distance≤8Å), our distance-based DL model can be used for contact prediction and has 3-4% better long-range prediction accuracy than the DL model directly trained from contact matrices.

In addition to predict C_β_-C_β_ distance distribution, we also train individual DL models to predict distance distribution for the following atom pairs: C_α_-C_α,_ C_α_-C_g_, C_g_-C_g_, and N-O. Here C_g_ represents the first CG atom in an amino acid. When CG does not exist, OG or SG is used. The predicted distance of these 5 atom pairs is used together to fold a protein, which on average is better than using the predicted C_β_-C_β_ distance alone. In addition, the predicted distance for C_α_-C_g_ and C_g_-C_g_ is also helpful for side chain packing.

### Predict secondary structure and torsion angles by 1D deep residual network

We employed a 1D deep ResNet of 19 convolutional layers to predict 3-state secondary structure and backbone torsion angles *ϕ* and *ψ* for each residue. The 1D ResNet has the same architecture as what we used for contact/distance prediction^10^ except the number of layers. Two types of input features are used: position specific scoring matrix (PSSM) generated by HHblits^36^ and primary sequence represented as a 20×L binary matrix where L is sequence length. For secondary structure, logistic regression is used in the last layer to predict the probability of 3 secondary structure types. However, for torsion angles we do not use logistic regression, but directly predict the angle distribution defined as follows.

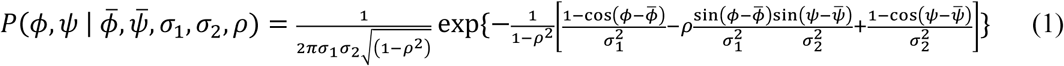

In Eqn.(1), 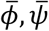 are the mean, σ_1_, σ_2_ are the variance and *ρ* is the correlation. That is, our deep ResNet outputs the mean and variance of the torsion angles associated with each residue. We use maximum-likelihood to train the network for secondary structure and angle prediction, i.e., maximizing the probability (defined by logistic regression or Eqn.(1)) of the observed properties of our training proteins. Note that the predicted mean and variance for angles is residue-specific, i.e., each residue has a different pair of predicted mean and variance. Our method for angle prediction is different from many existing ones, which usually discretize angles into some bins and then formulate the problem as a classification problem. The same set of training proteins (created in 2016) for distance prediction is used to train our DL models for secondary structure and angle prediction.

### Folding by predicted distance, secondary structure and torsion angles

Given a protein to be folded, we first predict its inter-atom distance matrix, secondary structure and backbone torsion angles, then convert the predicted information into CNS restraints and finally build its 3D models by CNS^30^, a software program for experimental protein structure determination. Given a matrix corresponding to the distance probability distribution for each atom type, we pick 7L (L is sequence length) of the residue pairs with the highest predicted likelihood (probability) having distance <15Å and assume the distance between these pairs is indeed <15Å. That is, we set their probability of having distance>16Å to be 0. From the predicted distance probability distribution, we may estimate the mean distance and standard deviation (denoted as *m* and s, respectively) of one atom pair, and then use *m-s* and *m+s* as its distance lower and upper bounds. We used the same method as CONFOLD^37^ to derive hydrogen bond restraints from predicted alpha helices. CONFOLD derived backbone torsion angles from predicted secondary structure, but we use the mean degree and variance predicted by our 1D deep ResNet as torsion angle restraints.

For each protein, we run CNS to generate 200 possible 3D decoys and then choose 5 with the least violation of distance restraints as the final models. CNS uses distance geometry to build 3D models from distance restraints and thus, can generate a 3D model within seconds. We generated multiple decoys for a protein since the CNS solution may not be globally optimal.

### The number of effective sequence homologs (Meff)

*Meff* measures the number of non-redundant (or effective) sequence homologs in an MSA, i.e., MSA depth. Here we use 70% sequence identity as cutoff to determine redundancy. Let *S*_*ij*_ be a binary variable indicating whether two protein sequences *i* and *j* are similar. *S*_*ij*_ is equal to 1 if and only if the sequence identity between *i* and *j* is >70%. For a protein *i*, let *S*_*i*_ denote the sum of *S*_*i1*_, *S*_*i2*, …,_ *S*_*i,n*_ where *n* is the number of proteins in the MSA. Then, *Meff* is calculated as the sum of *1/S*_*1*,_ *1/S*_*2*_, …,*1/S*_*n*_. Meff measures the difficulty of contact prediction especially for the DCA methods. The smaller Meff a target has, the harder it is for contact prediction.

### Performance metrics

We mainly use TMscore^38^ to evaluate the quality of a 3D model, which measures the similarity between a 3D model and its native structure (i.e., ground truth). It ranges from 0 to 1 and usually a 3D model is assumed to have a correct fold when it has TMscore≥0.5. Sometimes RMSD is also used to measure the deviation (in Å) of a 3D model from its native structure.

We measure the accuracy of predicted distance using 5 metrics: absolute error, relative error, precision, recall and *F1*. Since only predicted distance≤15Å is used as restraints to build 3D models, only the atom pairs with predicted distance≤15Å are considered. We define absolute error as the absolute difference between predicted distance and its native value, relative error as the absolute error normalized by the average of predicted distance and its native, precision as the percentage of atom pairs with predicted distance≤15Å that indeed have native distance≤15Å and recall as the percentage of atom pairs with native distance≤15Å that have predicted distance≤15Å. *F1* is calculated as *2×precision×recall/(precision+recall)*.

## Acknowledgement

This work is supported by National Institutes of Health grant R01GM089753 to JX and National Science Foundation grant DBI-1564955 to JX. The authors are also grateful to the support of Nvidia Inc. The funders had no role in study design, data collection and analysis, decision to publish, or preparation of the manuscript. The author is grateful to postdoc Dr. Sheng Wang, who helped generate multiple sequence alignment and input features for the CASP13 targets. The author is also grateful to Mr. Matthew Mcpartlon, who proofread this manuscript.

## Appendix

**Figure S1.**
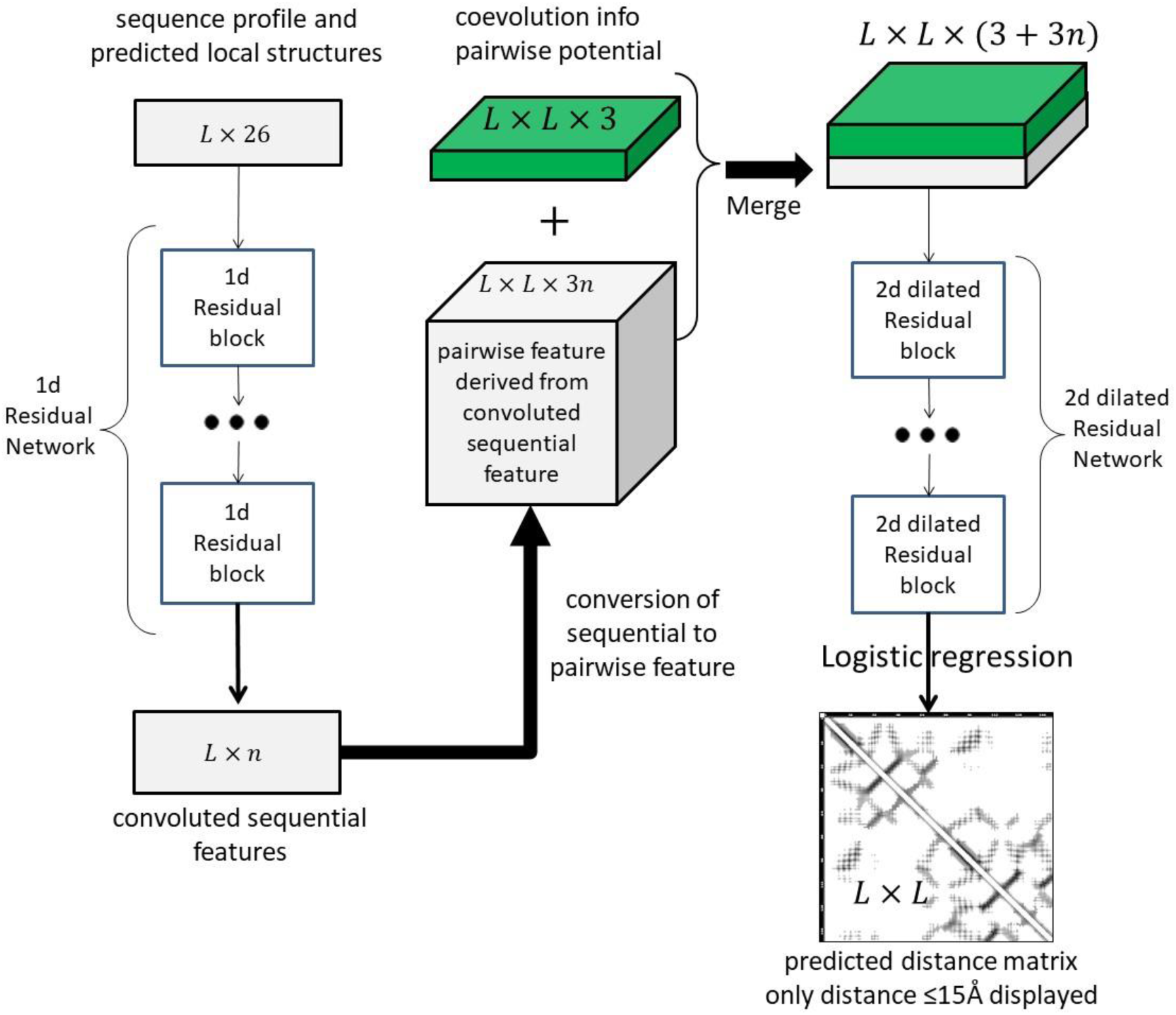
The overall deep network architecture for the prediction of protein distance matrix. The left column is a 1D deep residual neural network that transforms sequential features (e.g., sequence profile and predicted secondary structure). The right column is a 2D deep dilated residual neural network that transforms pairwise features. The middle column converts the convoluted sequential features to pairwise features and combine them with the original pairwise features. The picture is adapted from Figure 1 in the paper at https://journals.plos.org/ploscompbiol/article?id=10.1371/journal.pcbi.1005324.

